# Targets of sexual selection on the *Drosophila* wing

**DOI:** 10.1101/2024.10.21.619568

**Authors:** Stephen P. De Lisle, Anneli Branden, Ellen Paajarvi Vag

**Affiliations:** Karlstad University

**Keywords:** Sexual selection, geometric morphometrics, phenotypic selection, wing interference patterns

## Abstract

Identification of traits causally linked to fitness allows for direct tests of the adaptive value of traits. In the case of *Drosophila* and other insects, wing interference patterns – striking structural color variation generated by optical thin-film interference – have been implicated in mate choice and sexual selection. Yet, we expect wing interference patterns may covary with many aspects of wing morphology, such as size and shape. Here, we used a competitive mating assay in *Drosophila melanogaster* to assess the degree to which sexual selection acts on wing color independently of wing size and shape. We found that 16% of multivariate wing color, and 17% of corresponding estimated wing thickness, can be explained by wing size and shape. Projection pursuit regression identified three suites of traits – a linear combination of color variables, wing size, and a linear combination of wing shape variables – as the most important predictors of fitness. Analysis of the corresponding selection gradients revealed a combination of strong directional selection on wing size, combined with multivariate stabilizing selection on wing size and shape, with weak but significant directional selection on wing color. Our results suggest that sexual selection may act on a complex combination of wing color and morphology in flies.

## Introduction

Elaborate display traits are commonplace in nature and are often a clear outcome of sexual selection. In most cases, mating display traits are some of the most striking features of the organisms that express them, and as such their existence and importance long noted by naturalists (Darwin 1871, Endler 1992, Andersson 1994, Houde 1997). An intriguing exception to this lies in the Diptera, where striking wing coloration patterns have only been identified relatively recently as a potential mating display trait (Shevtsova et al. 2011, Katayama et al. 2014). Wing interference patterns are bold, iridescent color patterns clearly visible on otherwise-transparent fly wings under certain lighting conditions (Figure 1). These color patterns are formed by the phenomenon of optical thin-film interference, the same physical process that generates iridescent colors on soap bubbles and oil films (Shevtsova et al. 2011). These colors are formed by interference between light waves reflected off of the top surface of the wing and light waves that travel through the wing to be reflected off of the back surface; the combination of wing thickness and refractive index determines difference in speed between the two light waves and the resulting interference color generated (Newton 1704). Thus, wing interference patterns are the result of two structural features of fly wings; wing thickness and refractive index, the latter of which is expected to be relatively constant.

**Figure 1.**
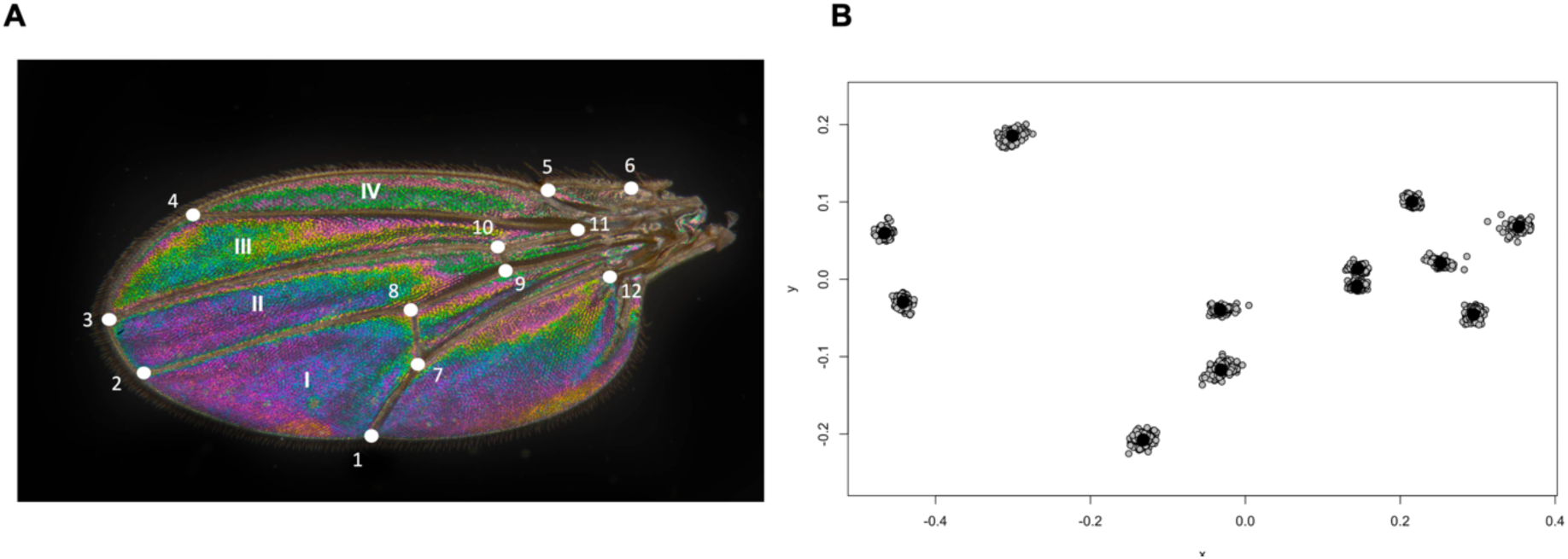
Image analysis of fly wings. Panel **A** shows wing interference color patterns produced under episcopic illumination by optical thin-film interference; wings are transparent when viewed under transmitted light. Also shown in **A** are the 12 landmarks used in analysis of wing shape as well as the four wing panels selected for measurement of rgb color. Panel **B** shows landmark variation after Procrustes rotation and scaling.

Recent work has highlighted wing interference patterns as a potential key trait in sexual selection and species divergence in flying insects. Diptera and other insect species are readily distinguished taxonomically on their wing interference patterns, which are often sexually dimorphic (Shevtsova et al. 2011, Hosseini et al. 2019, Butterworth et al. 2021, Takahashi 2022, Li et al. 2023). In fruit flies, *Drosophila,* wing interference patterns have been shown to be under sexual selection in males (Katayama et al. 2014), and wing interference patterns evolve in response to experimental manipulation of sexual selection (Hawkes et al. 2019). Consistent with the latter result, wing interference patterns are also observed to be heritable in different lineages of *D. melanogaster* (Katayama et al. 2014, Li 2014), and wings are known to be important for courtship in *Drosophila* (Greenspan and Ferveur 2000). All of the above evidence suggests that wing interference patterns are an important target of sexual selection in flies, and particularly in *D. melanogaster*.

Unlike some other elaborate display traits, wing interference colors are inevitably produced by transparent thin films, and thus are expected to be associated with such structures even if not a target of selection themselves. For example, in their comparative study of hoverflies, Li et al. (2023) found a strong relationship between wing size and interference reflectance spectra (and inferred wing thickness). Of past studies of *Drosophila*, only Li (2014) included some measure of wing morphology (wing length and width). Thus it is unclear the extent to which wing interference color is related to wing shape in *Drosophila,* which is noteworthy given the pervasive variation in wing shape maintained over deep time (Houle and Meyer 2015, Houle et al. 2017), and that wing size and shape differ across the sexes (Sztepanacz and Houle 2021). Here, we combine a mate choice assay in *Drosophila melanogaster* with detailed measurements of male wing interference color and multivariate wing morphology, in order to perform multivariate selection analysis (Lande and Arnold 1983, Svensson 2023) and identify the targets of sexual selection on the *Drosophila* wing. Our results indicate that while wing color is a direct target of sexual selection, selection appears to act on a complex combination of wing interference color, size, and shape.

## Methods

### Data collection

Each wild type male fly (LH_m_ background) was placed in a vial with two other unmated flies, a male and a female each homozygous for the *bw* brown eyed mutation but otherwise of similar genetic background. After 24 hours all flies were removed, resulting offspring allowed to develop, counted and scored for eye color; flies with wild type (red) eyes were sired by the focal male, giving a measure of binomially-distributed reproductive success in a competitive mating context, and thus a relevant component fitness measure for measuring sexual selection (Arnold and Wade 1984). That is, for each fly we had the count of offspring sired by the focal male as well as the offspring sired by the competitor male, and thus a count of “success” and “fail”. This is a standard approach for assessing competitive mating success in *Drosophila* (Chippindale et al. 2001); further details on the flies in this study can be found in (De Lisle 2024). We focused on the binomial measure of absolute fitness (counts of red-eyed offspring relative to brown-eyed offspring) in subsequent phenotypic selection analysis. Following this assay, all flies were frozen, thawed and left at room temperature. The right wing of each focal fly was then removed and photographed under a stereomicroscope (Olympus SZX-16) fitted with a 0.8x plan-apochromatic objective; the objective was aligned with the photographic optical path to minimize distortion and chromatic aberration. In case of excessive damage to the right wing, the left wing was instead photographed and the resulting image inverted digitally on the horizontal axis. Wings were placed under a cover slip on a microscope slide placed over an open aperture, and illuminated from above with a ring light to reveal interference patterns (see Figure 1). White balance on the camera (Nikon D850) was customized for the ring light illuminator, and all images were taken at identical exposure, illumination settings, and magnification. Otherwise-unmanipulated raw image files were downsized to 2000 x 3000 pixels for ease of analysis (noting that the native resolution of the camera exceeds the resolving power of the optical system) and saved as RGB jpeg files.

For each wing image, we placed 12 fixed landmarks by hand on vein intersections, using the same 12 landmarks that have been widely used to characterize variation in *Drosophila* wing shape (Houle and Meyer 2015, Houle et al. 2017). We then performed a generalized Procrustes analysis of these landmarks, followed by principal component analysis of the resulting geometric morphometric data to obtain 20 principle components describing multivariate wing shape. Our rotation (Table S1) was based on landmarks from a total of 232 flies, which included some females and some males for which we had no fitness data; all subsequent analyses beyond this initial rotation were performed only on the 170 males for which we had wing measurements and fitness data. We retained all 20 shape components (noting that three degrees of freedom are lost due to superimposition in the Procrustes analysis, and a further for centroid size) in subsequent analysis, as extensive past work in *D. melanogaster* suggests that all components harbor genetic variance (Houle et al. 2017). We also retained centroid size as a measure of overall wing size. We performed landmark placement, Procrustes analysis, and the principle component analysis using the *geomorph* package in R (Adams and Otárola-Castillo 2013).

We obtained information on wing interference color by using the color histogram tool in Fiji (Schindelin et al. 2012) to calculate mean values for red, green, and blue color channels for manual selections of regions of each wing. Although other studies have focused instead on the color triplet of hue, saturation, and brightness, we focus on RGB color because this is the data recorded by a Bayer camera sensor. We focused on four major wing panels that are clearly demarcated by vein intersections and capture most of the wing surface area (Figure 1A), resulting in a total of 12 color traits for each wing. We performed a principal component analysis on the covariance matrix to obtain 12 orthogonal color variables. We also calculated mean RGB values for the entire manually-selected wing, and present analysis of this simplified color data in the supplement.

We used an inverse colorimetry approach to infer the average thickness of each of the four wing panels for each fly. In our images, thin film interference generates reflectance spectra that are recorded by a Bayer sensor in the digital camera. A direct estimate of the reflectance spectra (e.g., Li et al. 2023) would allow direct calculation of thickness of a film with known refractive index. In our case, however, our recordings of the spectra were not direct; rather, the camera records light intensities in three color channels (Stevens et al. 2007). While there are multiple reflectance spectra that could generate any given RGB triplet, it is possible to generate a “most likely” estimated reflectance spectrum by taking the centroid of all possible spectra that could generate a given RGB triplet (Davis 2019, 2024). We performed such an analysis using the invert() function in the coloSpec package in R (Davis 2024). We inferred spectra from the mean RGB value separately for each wing panel, over a wavelength range of 450-650 nanometers (corresponding to the peak transmittance range of the UV-IR cutoff filter of most digital camera sensors), and assuming a color temperature of 5600K corresponding to the Schott ring light illumination used. Inferred reflectance spectra are illustrated in Figure S1. We then used nonlinear least squares to fit equation 8 from (https://farbeinf.de/pdf/soapfilmcalc.pdf) to each inferred reflectance spectrum, estimating two parameters: a constant, *C,* and *s*, where *s* is parameter that is related to the product of wing thickness (*d*) and refractive index of the wing (*n*) by *s = 2dn.* We were then able to calculate wing thickness in nanometers assuming a refractive index of chitin of *n* = 1.53. All nonlinear models converged and yielded estimated thickness values in the expected order of magnitude (approximately several hundred nanometers), however we emphasize that our inferences of wing thickness should be viewed with extreme caution; without spectrometer measures, we present them primarily as a proof of concept. We excluded several outliers from downstream analysis as they had extreme leverage in regressions with fitness.

### Data analysis

For each male fly we thus obtained data on reproductive success, multivariate wing morphology (20 shape PCs plus centroid size) and multivariate wing interference color (12 color PCs) for a total of 33 traits. First, we used redundancy analysis (RDA) to assess the relationship between the color traits and the morphology traits, specifically the degree to which variation in multivariate color can be explained by variation in wing morphology. We fitted the RDA using the vegan package in R, and used permutation ANOVA (at defaults of rda() in vegan) to assess statistical significance of the model and each RDA axis. We found statistical support for the first RDA axis, which represents a linear combination of morphological traits that predicts the most variation in wing color. We used a Bayesian multi-response mixed effects model (Hadfield 2010) to assess differences in inferred wing thickness between panels, presenting 95% highest posterior density (HPD) intervals for pairwise contrasts between panels.

For our phenotypic selection analysis, we used projection pursuit regression to reduce dimensionality and identify which of the 33 traits are potential targets of selection. We first fitted a projection pursuit regression using all 33 traits as predictors, and the logit of reproductive success, 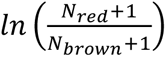, as the response and the total number of offspring from the mating assay as a weight term (noting that it is not readily possible to fit a binomial or quasibinomial projection pursuit using currently available functions in R). We calculated standard errors and p-values for the loadings on the first projection pursuit axis using a bootstrap (n = 500 samples); this procedure identified three traits that loaded significantly on this axis, color PC1, centroid size, and shape PC 5. We then refitted the projection pursuit regression using only these three traits, since inclusion of traits unrelated to fitness can lead to inaccurate estimates of selection (Sztepanacz and Houle 2024).

We then performed a phenotypic selection analysis using this reduced set of three traits identified by the projection pursuit regression: color PC1, centroid size, and shape PC 5. We then fit two generalized linear models for estimation of selection gradients, one with just linear terms for these three traits (noting that linear gradients should be obtained from a regression without nonlinear terms; Lande and Arnold 1983), and a second quadratic GLM with all linear and nonlinear terms to estimate nonlinear selection (Phillips and Arnold 1989). We modelled absolute fitness, which was the joint number of successes (red-eyed offspring) and failures (brown-eyed offspring), as the response (performed using cbind() to couple these counts on the left side of the model formula in glm() in R). We assumed a quasibinomial error distribution after encountering over dispersion when fitting a binomial model; although a pure Bernoulli (0 or 1) outcome cannot be over dispersed, counts of successes and fails (which constitute multiple Bernoulli trials) can be and this was the case with our data. We then computed standardized selection gradients from our GLM coefficients following the approach outlined by Janzen and Stern (1998), and weighting by the inverse of mean absolute fitness (red eyed/total offspring) calculated across the experiment (De Lisle and Svensson 2017). In order to obtain standard errors for these resulting gradients we performed a bootstrap wherein we re-fit the GLM in each bootstrap sample (n = 1000), calculating the gradients as described above, and took the standard deviation of the resulting sampling distribution as an estimate of the standard error of the gradient. We multiplied our estimates of quadratic selection by two to make them interpretable nonlinear gradients (Stinchcombe et al. 2008).

Our analysis of nonlinear selection yielded a 3-dimensional γ matrix of nonlinear selection gradients. In order to aid interpretation of this matrix we performed a canonical rotation (eigen analysis) to identify orthogonal axes of stabilizing or disruptive selection. This decomposition yielded 3 eigenvalues and corresponding eigenvectors, and in order to assess statistical significance of the eigenvalues we used a modification of the Reynolds et al. (Reynolds et al. 2010) permutation procedure (Chenoweth et al. 2012, De Lisle and Rowe 2015). This permutation (n = 1000) involved permuting fitness along the eigenvectors identified in the original rotation of γ, fitting a quadratic GLM with these eigenvectors as predictors of fitness, and comparing *F* statistics from nonlinear terms to the corresponding *F* statistics computed the same way from the non-permuted data; the proportion of permutation *F* statistics greater than the original observed gives an estimate of the p-value for the corresponding eigenvalue of γ.

We repeated the above analysis (projection pursuit regression + GLMs with reduced trait number) using inferred thickness of the four wing panels instead of the 12 color traits. All traits were mean centered and variance-standardized for fitting regressions (both projection pursuit and GLMs). Data and code are available as supplemental material and at Zenodo LINK REDACTED.

## Results

We obtained complete data on wing color, morphology, and fitness for a total 170 male flies, with fitness data representing eye color score for 7875 total offspring. Wing color traits were generally highly correlated both within and across wing panels (Figure 2). A redundancy analysis (RDA) revealed that 16% of the variation in multivariate wing color can be explained by (is redundant with) the 21 wing size and shape variables (permutation ANOVA *F*_21,148_ = 1.34, *P* = 0.0002). Permutation ANOVA revealed statistical support for the first RDA axis (*F*_1,157_ = 5.92, *P* = 0.016), but not lower axes (all P ≥ 0.097). Relationships between color traits and wing morphology are illustrated in Figure 2. In a simplified analysis of mean red, green, and blue values across the entire wing, we found concordant results (16.7% of mean color variation explained by wing morphology; permutation ANOVA *F*_21,148_ = 1.41, *P* = 0.062). We also found similar results in an RDA predicting wing depth from morphology (17.4% of wing thickness variation explained by wing morphology; permutation ANOVA *F*_21,135_ = 1.35, *P* = 0.096). Our inference of wing thickness indicate wing thickness of approximately 300 nanometers (Figure 3), generally consistent with past work (Katayama et al. 2014, Hadjaje et al. 2024) as well as our own observations that wings of LH_m_ *Drosophila* are just thinner than the usable axial resolution limit of an oil immersion light microscope. There were significant differences among wing panels, with panels 2 and 4 inferred to have greater thickness on average than panels 1 and 3 (Figure 3; Table S7). Depth measures were strongly correlated across wing panels, with positive correlation between panel 1 and 2 depth, 3 and 4 depth, and negative correlations across panels 1,2 and 3,4 (Figure S2).

**Figure 2.**
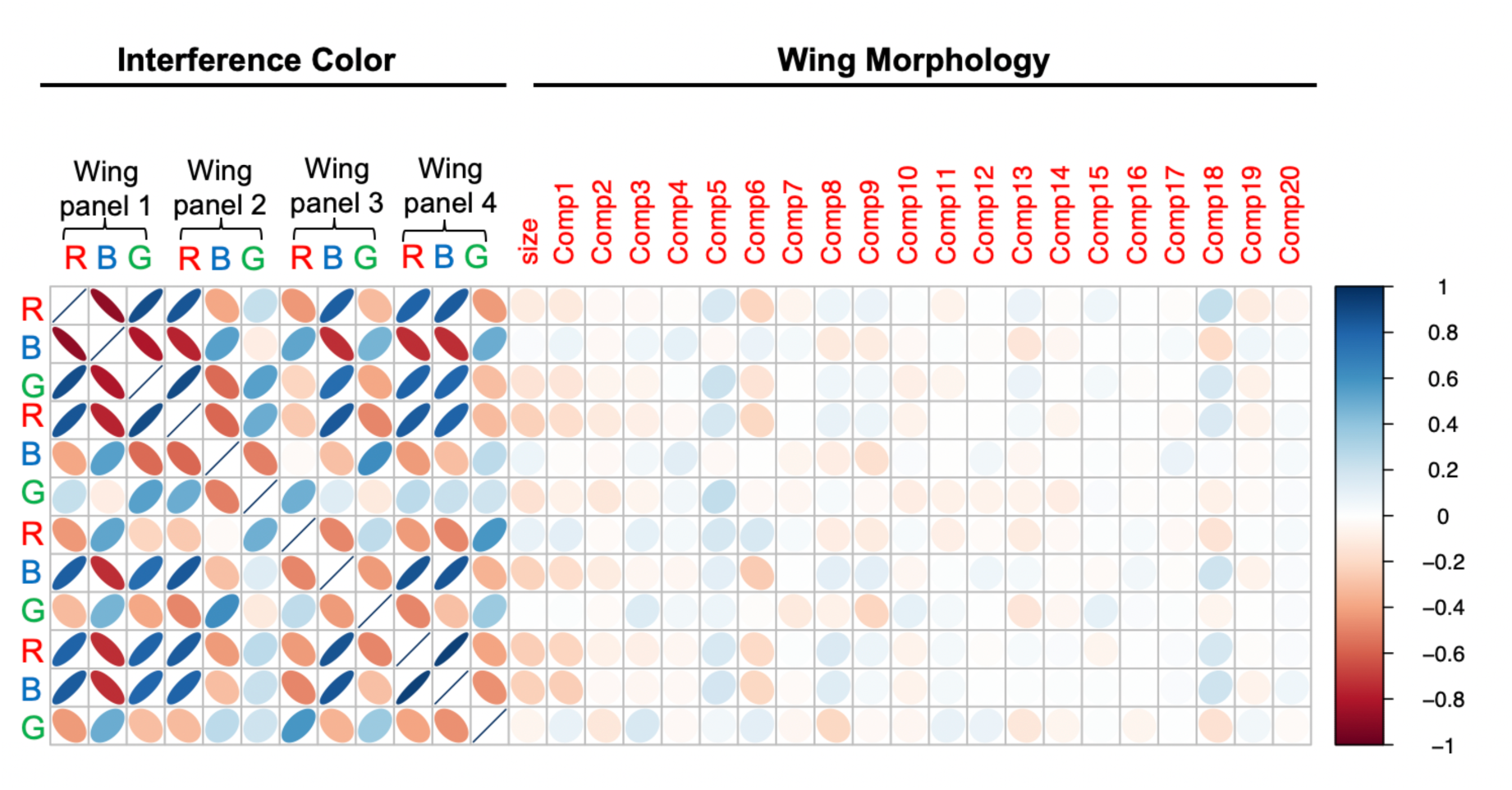
Correlations between wing color and morphology. Shown are the correlations between RGB color values for each wing panel, and the correlation between each of these color traits and the 21 morphological traits from the geometric morphometric analysis. Although correlations between color traits and morphology were far weaker than correlations between color traits, a redundancy analysis revealed a significant (16%) fraction of color variation can be explained by wing size and shape (see text).

**Figure 3.**
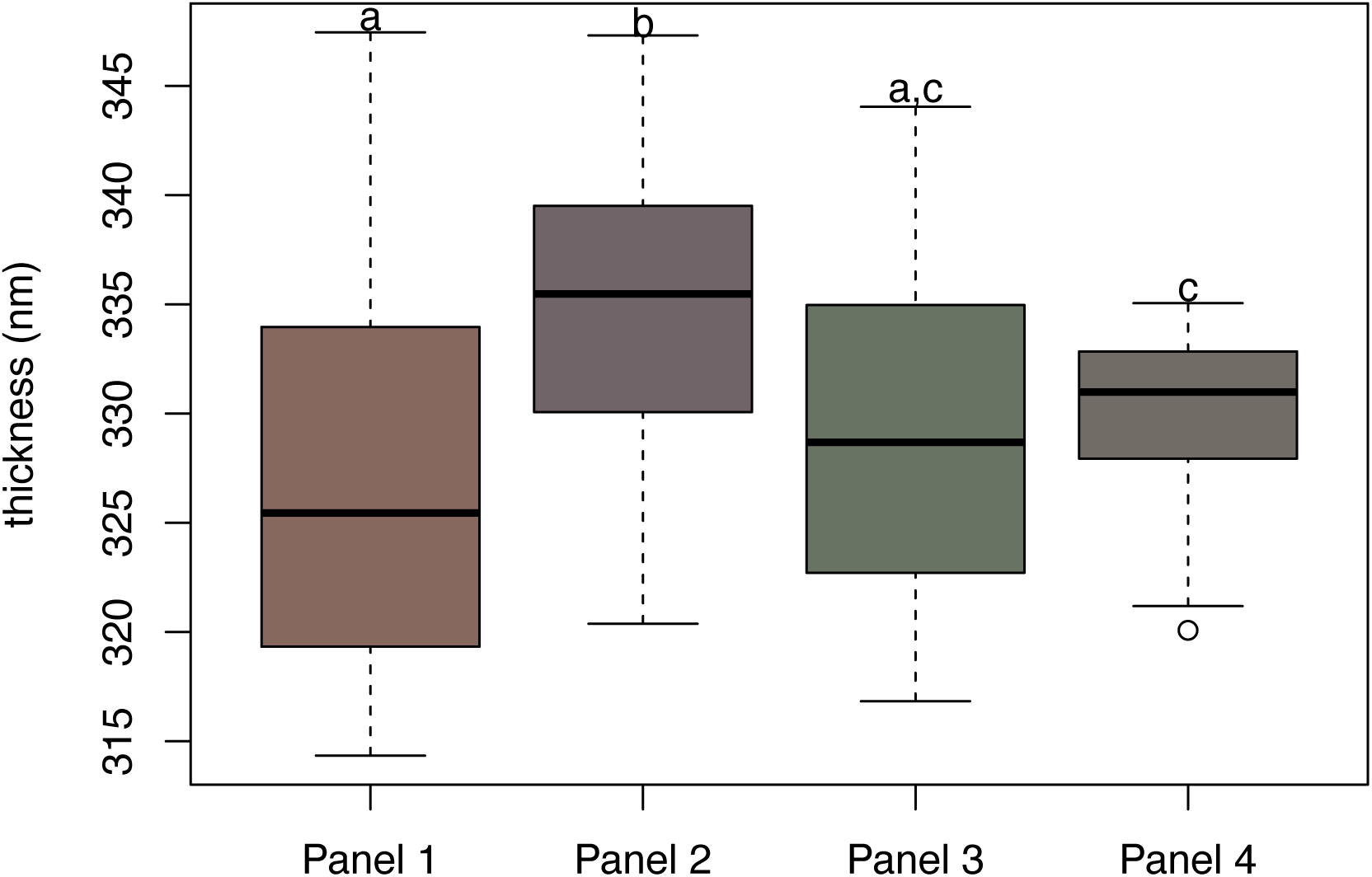
Inferred wing thickness across the four measured wing panels. Each box and whisker plot is colored according to the mean RGB triplet for that panel. Letters denote statistical significance of pairwise contrasts from a multiresponse mixed model. See text for details on inference of thickness.

**Figure 4.**
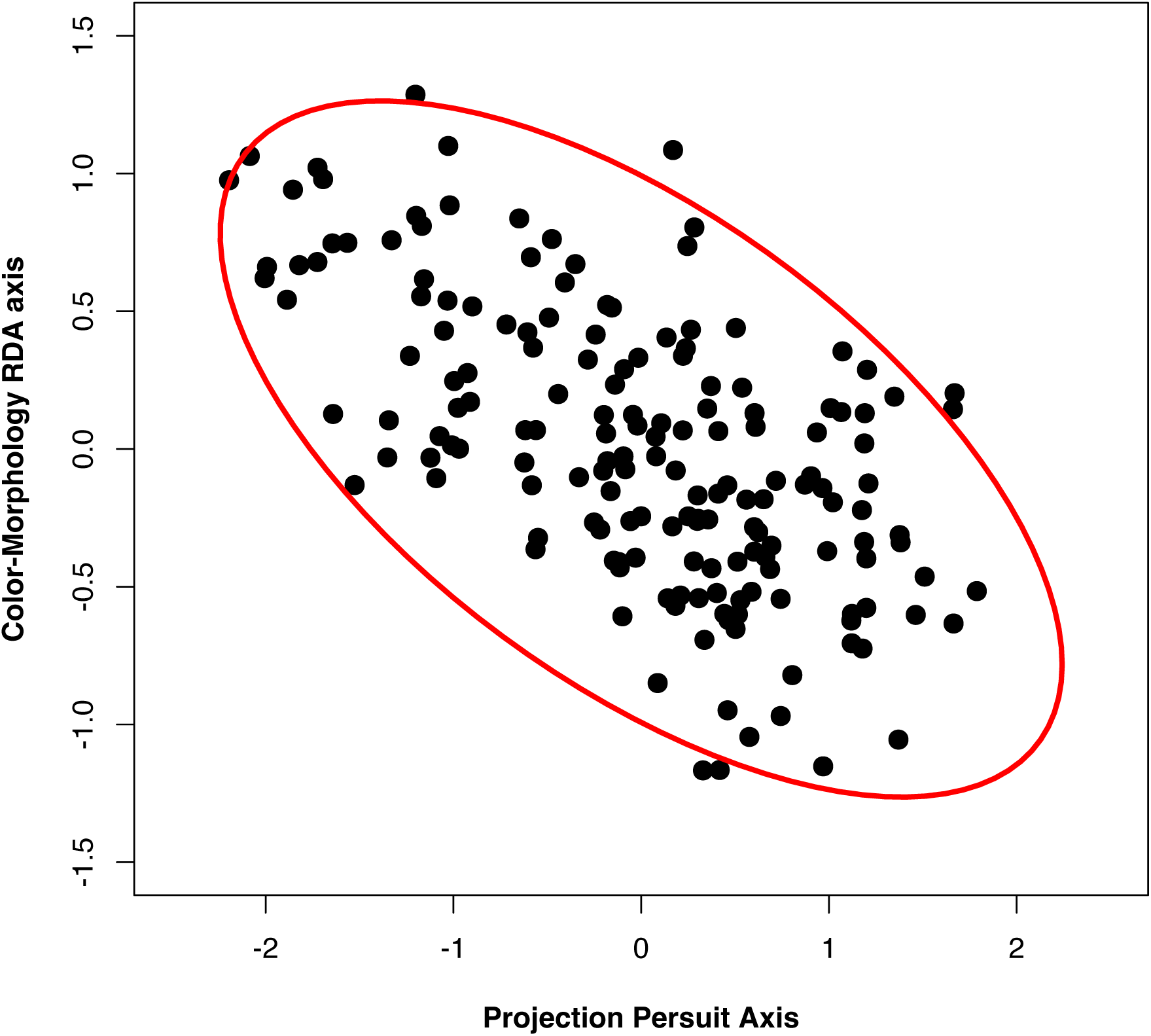
Correlation between redundancy (RDA) axis and the axis of strongest selection identified by projection pursuit regression. The RDA axis represents the linear combination of traits that is redundant between WIP color and morphology (size and shape); thus, this combination of traits is closely aligned with selection. Shown is the 95% confidence ellipse for the covariance between the two traits.

Projection pursuit regression identified 3 traits as the most important in explaining variation in fitness: PC 1 of the color traits (coefficient from reduced model: 0.43, bootstrap SE = 0.16), centroid size (coefficient from reduced model: 0.83, bootstrap SE = 0.11), and PC 5 of wing shape (coefficient from reduced model: -0.35, bootstrap SE = 0.22). This combination of traits was strongly correlated with the RDA axis (r = -0.62. t_168_ = -10.2, P >0.0001; Figure 3); that is, the linear combination of traits explaining the most variation in fitness was closely related to the combination of traits that is redundant between color and morphology. Examining the loadings on color PC1 (Table S2) reveals that color variation associated with wing panels 1 and 4 contribute most to this dimension (square root of sum of squared elements: 0.74 and 0.45 respectively), followed by panels 2 and 3 (square root of sum of squared elements: 0.38 and 0.29 respectively). Performing these calculations by color channel indicates a rank order importance for red, blue, and green (0.68, 0.57, 0.44, respectively). Loadings on shape PC 5 (Table S1) indicate that this axis is most strongly associated with variation in the horizontal dimension (square root of sum of squared elements: 0.82 versus 0.56) and landmarks 4 and 7 (Figure 1; square root of sum of squared elements: 0.42 and 0.43).

Examining the fitness surface along this projection pursuit axis reveals a fitness surface characterized by a combination of directional and stabilizing selection (Figure 5A). When fitting GLMs to the original traits contributing to this projection pursuit axis, we found significant directional selection on centroid size (*t*_166_ = 4.8, P < 0.0001) and wing color PC1 (*t*_166_ = 2, P = 0.045; Table 1). Fitting a quadratic GLM revealed significant stabilizing selection on centroid size (*t*_160_ = -2.5, P = 0.013; Table 2) and a trend for stabilizing selection on wing shape PC 5 (*t*_160_ = -1.8, P = 0.069; Table 2). Standardized linear selection gradients indicate that the estimated strength of directional selection on centroid size (*β* = 0.165), was over twice as strong as selection on either of the other traits (Table 3). Differences among traits in the strength of nonlinear selection was less pronounced (Table 3), although canonical rotation of γ revealed a single significant eigenvalue (estimate = -0.166) representing stabilizing selection on a combination of traits dominated by centroid size, and to a lesser degree wing shape (Table 4). Also notable in this rotation is the trend towards significance (P = 0.08) for the second smallest eigenvalue (estimate = -0.079), representing stabilizing selection on a combination of traits dominated by wing shape. Thus this analysis reveals stabilizing selection acting predominantly on a combination of centroid size and PC 5 of wing shape. The fitness surface for the three traits is plotted in Figure 5A,B.

**Figure 5.**
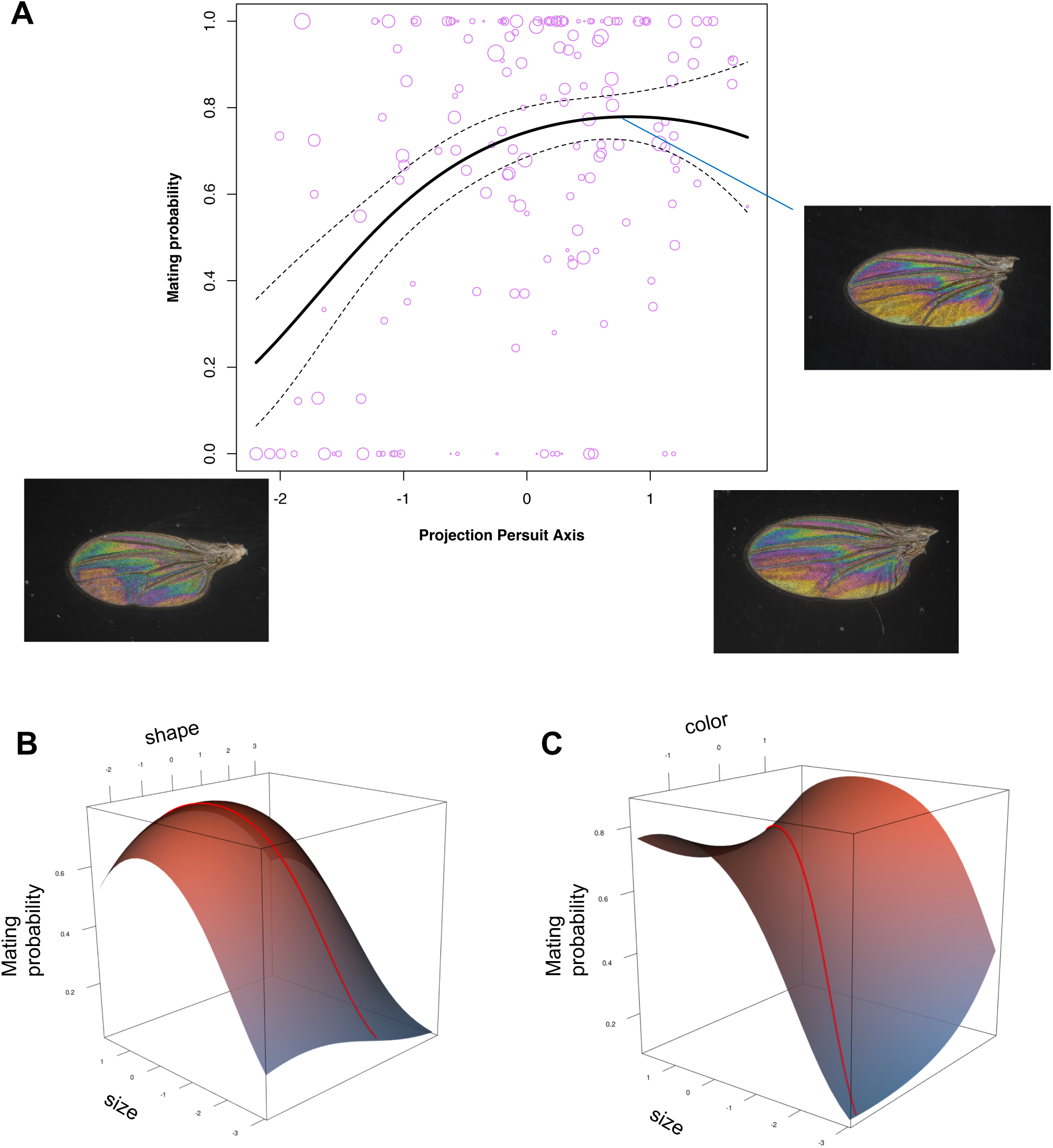
Fitness surface along projection pursuit axis. Panel **A** shows the relationship between mating success and multivariate wing color along the axis recovered from a projection pursuit regression with wing color, size, and morphological shape as predictors; this represents the direction in trait most associated with fitness and is characterized by a combination of directional and stabalizing selection. Line shows predicted values and 95% CI from a quadratic quasibinomial GLM fitted to the projection pursuit axis scores. Inset panels show wing images from flies with the lowest and highest score on the projection pursuit axis, as well as the individual with the highest predicted fitness from the GLM. Panels **B** and **C** shows the relationships between mating success and the three traits that contribute to the projection pursuit axis; color PC 1, centroid size, and shape PC 5, from quadratic quasibinomial GLMs. The red line illustrates the projection pursuit axis.

**Table 1.** GLM for linear selection analysis of wing color, size, and shape.

|  | **Estimate** | **SE** | **t** | **P** |
| --- | --- | --- | --- | --- |
| intercept | 0.8353 | 0.118 | 7.111 | 3.27E-11 |
| color PC1 | 0.2437 | 0.121 | 2.014 | 0.0456 |
| centroid size | 0.5432 | 0.113 | 4.801 | 3.50E-06 |
| shape PC 5 | -0.1444 | 0.123 | -1.178 | 0.2405 |

Residual deviance: 4108.2 on 166 degrees of freedom
dispersion parameter: 21.05905

**Table 2.** GLM for nonlinear selection analysis of wing color, size, and shape.

|  | Estimate | SE | t | P |
| --- | --- | --- | --- | --- |
| intercept | 0.97922 | 0.2293 | 4.27 | 3.34E-05 |
| color PC1 | 0.24457 | 0.1281 | 1.909 | 0.0581 |
| centroid size | 0.30787 | 0.1412 | 2.18 | 0.0307 |
| shape PC 5 | -0.16054 | 0.1221 | -1.315 | 0.1904 |
| color PC1 ^2 | 0.23176 | 0.1591 | 1.457 | 0.1472 |
| centroid size ^2 | -0.25934 | 0.1036 | -2.503 | 0.0133 |
| shape PC 5 ^2 | -0.14416 | 0.0789 | -1.826 | 0.0697 |
| color PC1*centroid size | -0.09556 | 0.1409 | -0.678 | 0.4986 |
| color PC1*shape PC 5 | 0.11026 | 0.1375 | 0.802 | 0.4238 |
| centroid size*shape PC 5 | 0.08473 | 0.1384 | 0.612 | 0.5414 |

Residual deviance: 3854.5 on 160 degrees of freedom
dispersion parameter: 21.04012

**Table 3.**
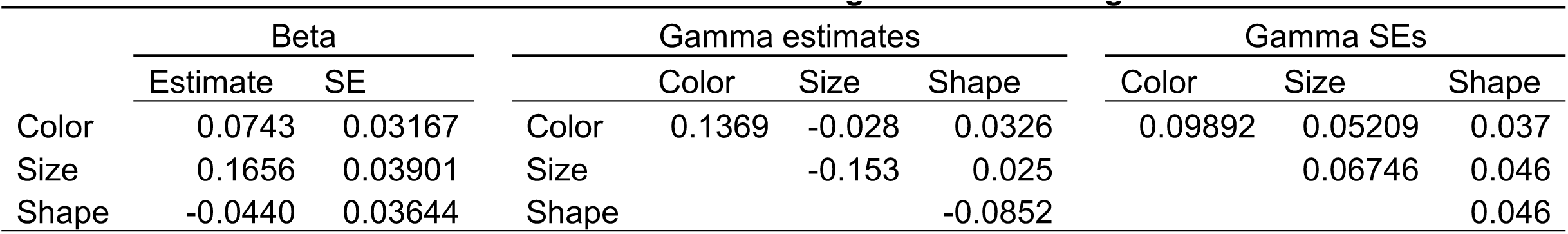
Standardized selection gradients for wing traits.

**Table 4.** Canonical rotation of γ.

|  | Eigenvalues |  |  |
| --- | --- | --- | --- |
|  | 0.1436 | -0.0791 | **-0.1660** |
|  | P = 0.164 | P = 0.083 | **P = 0.008** |
|  | Eigenvectors |  |  |
| Color | 0.9878 | -0.0947 | -0.1233 |
| Size | -0.0828 | 0.3505 | -0.9329 |
| Shape | 0.1316 | 0.9318 | 0.3384 |

Repeating our selection analysis but focusing on inferred wing thickness of the four measured panels instead of color estimates resulted in essentially the same conclusions. We found that a projection pursuit regression (Table S4) indicated three traits contributing most to fitness variation: panel 4 thickness, centroid size, and a trend for shape PC 5. Retaining these traits in a phenotypic selection analysis revealed evidence of positive directional selection on panel 4 thickness (*t*_153_ = 2.3, P = 0.0225; Table S5; *β* = 0.087, SE = 0.036). This analysis is thus consistent with the analysis of color variation in that we find evidence of directional selection limited to specific areas of the wing. We found no evidence of selection on the average color values measured across the entire wing (Table S6).

## Discussion

Using a competitive mating assay combined with morphometric analysis, we have shown that a complex combination of wing interference color and multivariate morphology appear to be targets of phenotypic sexual selection in *Drosophila melanogaster*. While directional sexual selection acts on features of multivariate wing color, the combination of wing traits that explains most variation in fitness is closely associated with the combination of traits that describes redundant variation between color and size/shape. Directional selection was over twice as strong on wing size compared to wing color, and nonlinear selection was characterized by stabilizing selection on wing size and shape. However, directional selection on color traits does remain after accounting for variation in wing size and shape, suggesting that wing color or thickness may indeed play a role in sexual selection. A major conclusion from our study is that interpreting fitness effects, or evolutionary response, of wing interference colors may be difficult without other measures of wing phenotype, although our results generally support past conclusions (Katayama et al. 2014, Li 2014, Hawkes et al. 2019) that wing interference colors may indeed be a target of sexual selection.

That said, our analysis reveals evidence of positive directional sexual selection on wing color acting simultaneously with stabilizing/directional selection on a combination of wing size and shape. Although selection (directional and nonlinear) was by far strongest on wing size, we found that selection also acted on combinations of shape and color variables. Selection on wing shape was strongest on the horizontal axis, nearly 50% greater than on the vertical axis, while selection color variation was strongest for the red color channel on panels 1 and 4. Flies with large, elongated wings with yellow WIPs obtained the highest fitness in our competitive mating assay, as illustrated in inset panels in Figure 5.

Although our approach to measuring reproductive fitness is a relatively standard method in *Drosophila*, it differed from behavioral mating assays of past studies of selection on wing interference. For example, other studies have recorded time to mating in no-choice assays as a measure of reproductive fitness, including manipulation of the background conditions that presumably affect the visibility of wing interference patterns (Katayama et al. 2014, Li 2014). While these past studies did not analyze wing shape variation and its fitness effects in the manner taken here, they did manipulate the selective environment to demonstrate that selection on these display traits indeed varies with the visual environment, as expected if they are a direct target of selection. This is in contrast to our methods, where we performed an assay of competitive reproductive success by scoring phenotypes of offspring resulting from 24 hours of potential mating, and while we measured traits expected to be correlated with wing color (i.e., performed a multivariate selection analysis; Lande and Arnold 1983, Svensson 2023), we made no effort to make wing interference patterns visible beyond what is typically observed in a standard fly vial under ambient light. Despite this different approach, we none the less recovered evidence of sexual selection on interference patterns, although we do not find pervasive evidence of stabilizing selection on this trait, as observed for wing hue by Katayama et al. (2014). We found evidence of strong stabilizing selection acted through a combination of wing size and shape variation that correlate with wing color.

We found that a small but significant fraction of interference color variation was redundant with wing size and shape. This redundancy was surprisingly low given that wing interference color is a function of wing thickness, which we may expect to be related to wing size and shape (Weber 1990). One explanation is that wing microstructure (Shevtsova et al. 2011) such as hair density contributes to wing thickness independently of wing size and shape. Despite this somewhat low redundancy of variation, we found that the combination of traits contributing most to this redundancy is strongly correlated with the combination of traits that explains most fitness variation; thus selection is largely acting on a combination of correlated wing color and morphological traits. This result can be understood in plain terms by realizing that wing size, itself correlated with many other wing traits, is under strong directional + stabilizing selection.

Using inverse colorimetry to infer wing thickness from images of interference color, we found some evidence for positive directional selection on wing thickness for one wing panel. This is consistent with the idea that wing color may correlate with quality or condition (Rowe and Houle 1996, Bonduriansky 2007), as has been previously suggested could be the case for *Drosophila* wing interference patterns (Katayama et al. 2014). However, we also found that wing thickness measures correlated in complex ways with each other, suggesting that the relationship between wing thickness and condition may not be straightforward. One explanation could be if wing hair density contributes to interference color, as it likely does (Shevtsova et al. 2011). Regardless, our analysis of wing thickness should be viewed as preliminary; without a calibrated interferometer or some other independent measure of wing thickness (e.g., Li et al. 2023), our measures of thickness primarily represent a proof of concept. Notably however our conclusions from the analysis of wing thickness were similar to our conclusions from the analysis of the raw color principal components, where color variation associated with panels 1 and 4 where under strongest directional selection.

One intriguing aspect of our results is our finding of stabilizing selection on wing size and particularly shape, which we largely expect to be predominant in a lab-adapted population. Although simple in structure, *Drosophila* wings harbor patterns of multivariate variation that has been conserved for millions of years (Houle et al. 2017). Relatively little is known about the contribution of selection to shaping this and other examples of deeply-conserved variation (McGlothlin et al. 2018, Henry and Stinchcombe 2023, De Lisle et al. 2024), yet it seems likely that coupling measures of multivariate fitness surfaces (Hohenlohe and Arnold 2008, Punzalan and Rowe 2016) with estimates of standing variation may be fruitful in illuminating how selection may maintain patterns of multivariate variation. Although our study lacked power to infer multivariate nonlinear selection on a full set of wing shape traits, future work that bridges larger-scale laboratory measures of selection with comparative biology may provide insight into how selective forces acting on complex traits can generate conservation of variation in deep time.

