## supplemental material for "Targets of sexual selection on the *Drosophila* wing"

**Supplementary Material for “Targets of sexual selection on the *Drosophila* wing”**

**Figures**

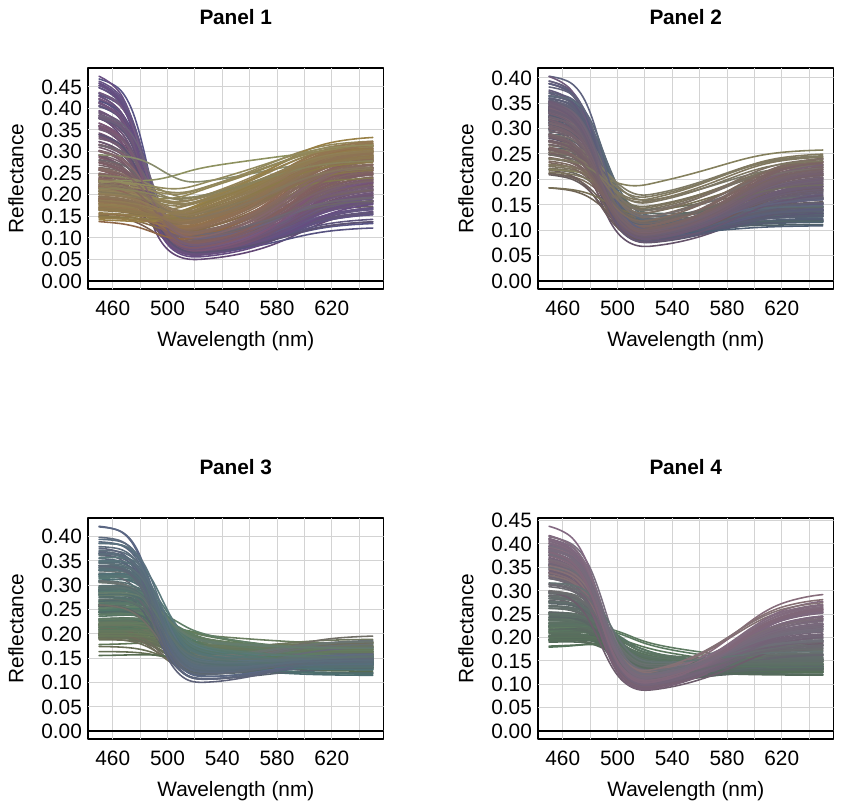

**Figure S1. Inferred reflectance spectra for each wing panel.** Shown is inferred relationship between the wavelength and reflectance inferred from RGB triplets for each wing panel. Individual curves correspond to individual flies. See text for methodological details.

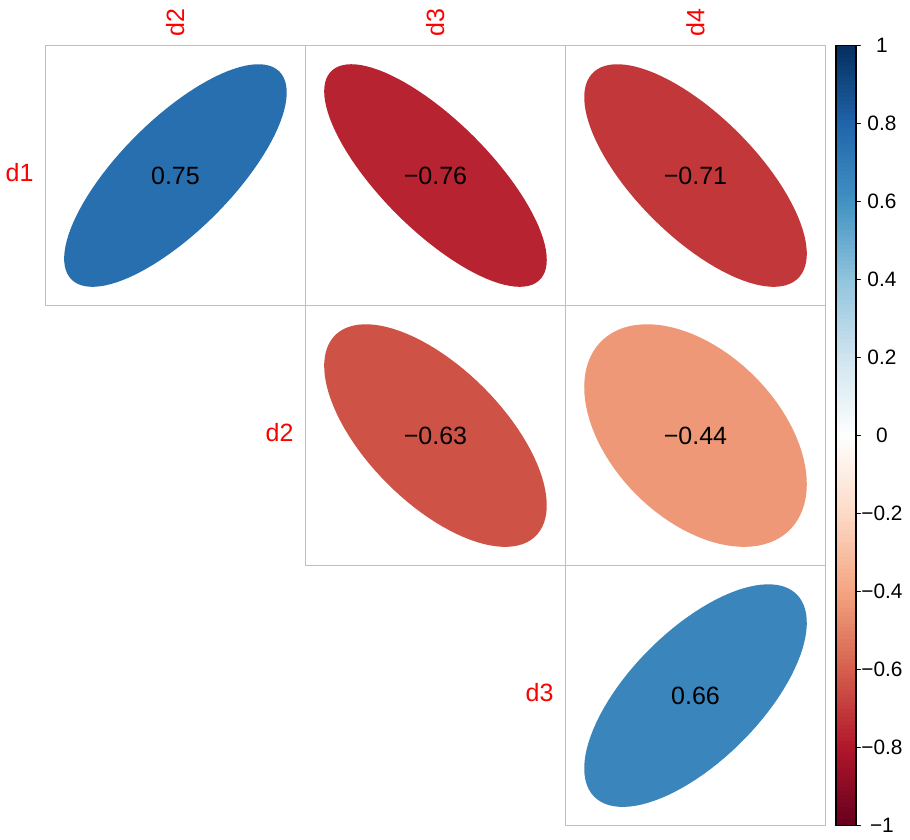

**Figure S2. Phenotypic correlation matrix for wing depth.** Shown are the phenotypic correlations for inferred wing depth across the four wing panels measured. All correlations were statistically significant.

**Tables**

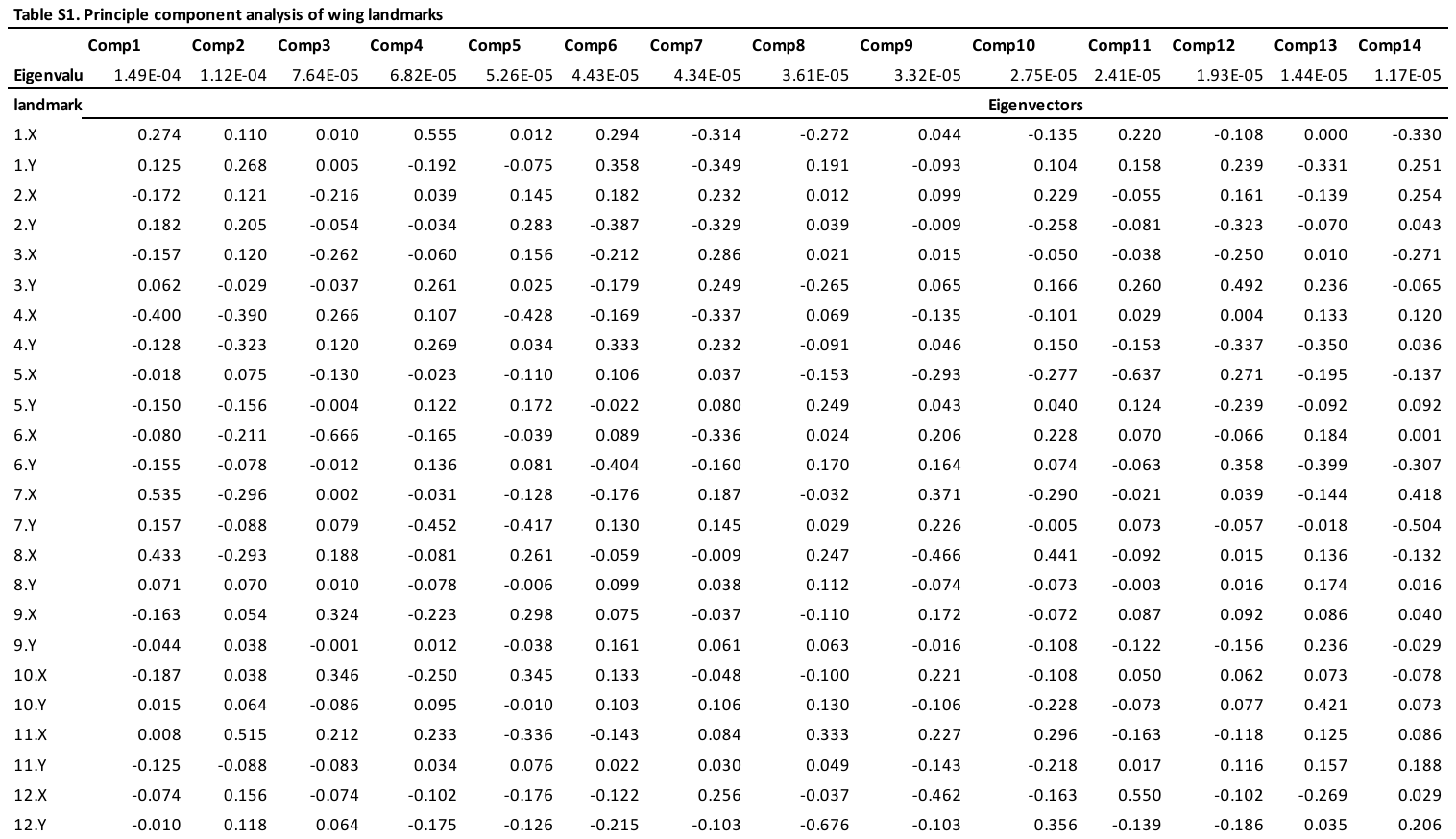

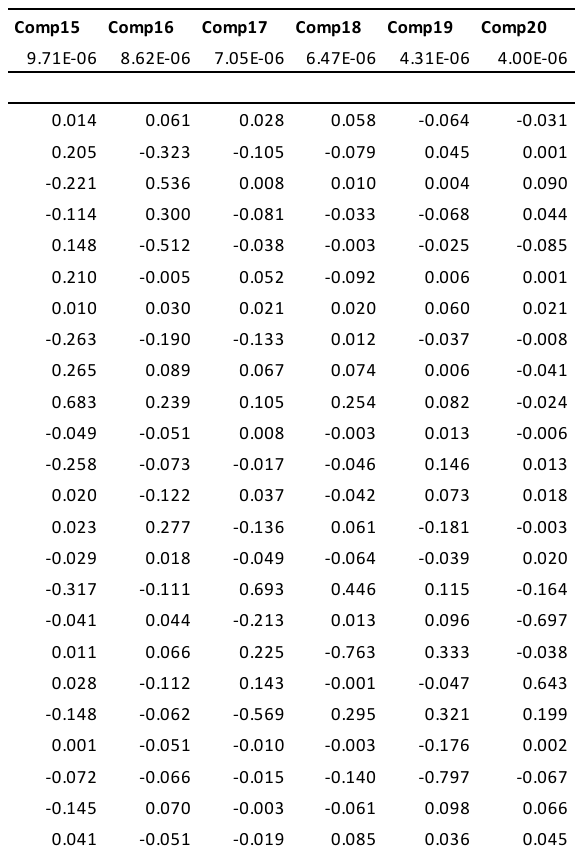

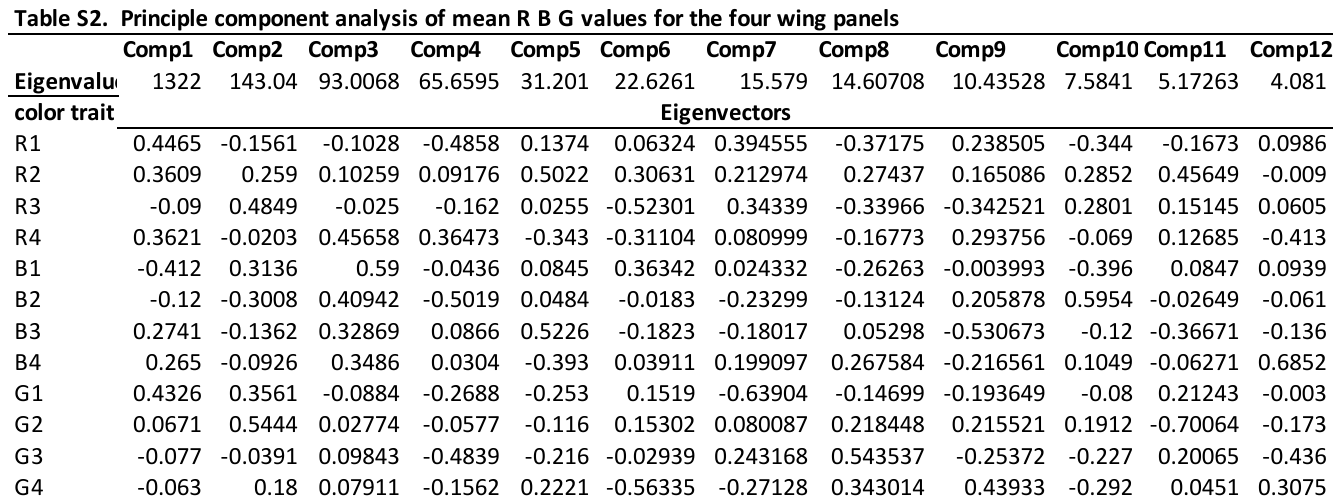

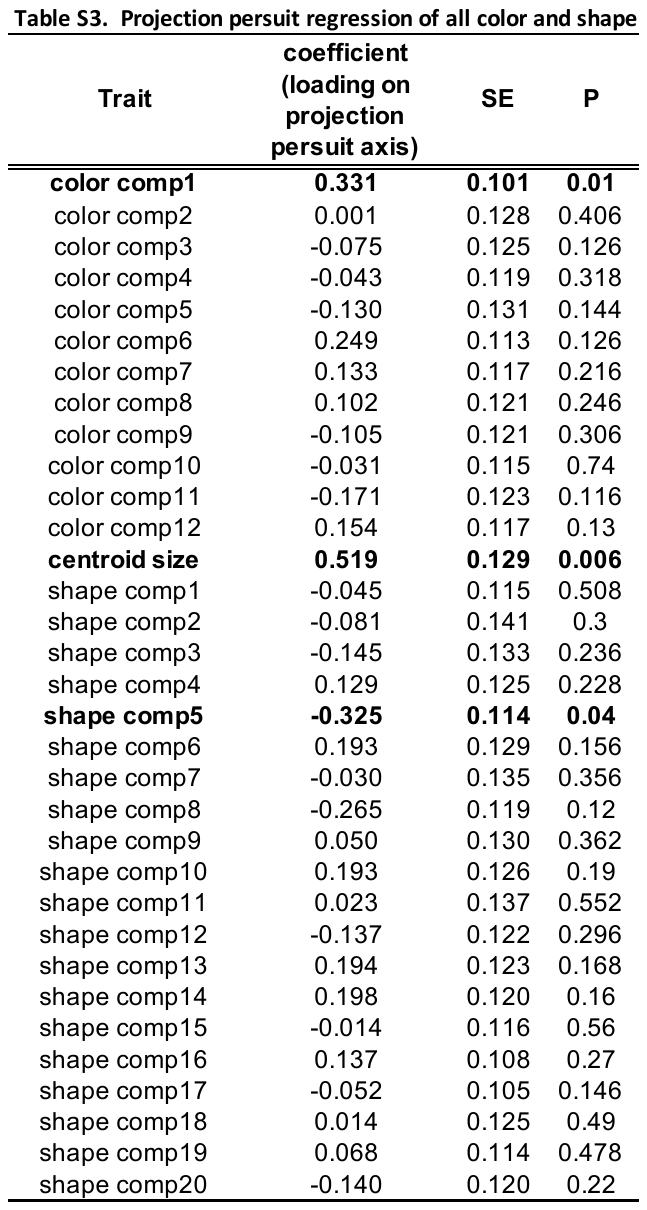

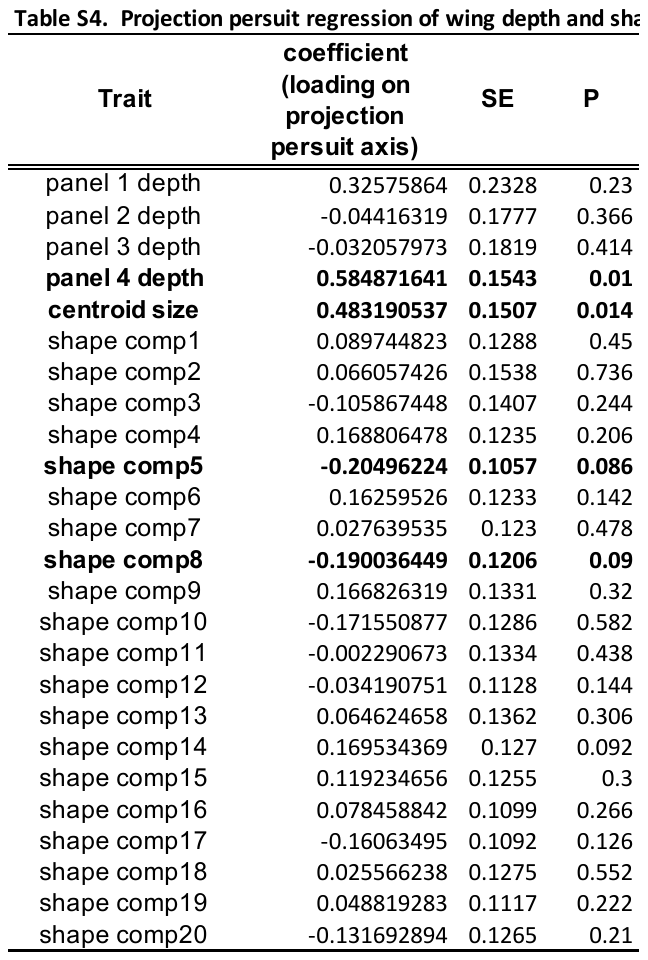

| **Table S5. GLM for linear selection analysis of inferred wing thickness, size, and shape** | | | | |
| --- | --- | --- | --- | --- |
|  | **Estimate** | **SE** | **t** | **P** |
| intercept | 0.8146 | 0.124 | 6.567 | 7.54E-10 |
| Panel 4 thickness | 0.2852 | 0.1237 | 2.305 | 0.0225 |
| centroid size | 0.5483 | 0.1177 | 4.66 | 6.84E-06 |
| shape PC 5 | -0.1335 | 0.1286 | -1.038 | 0.3007 |
| Residual deviance: 3868.8 on 153 degrees of freedom | | | | |
| dispersion parameter:21.74743 | |  |  |  |

| **Table S6. GLM for mean whole-wing color** | | | | | | | | |
| --- | --- | --- | --- | --- | --- | --- | --- | --- |
| **Trait** | | **Estimate** | | **SE** | | **t** | | **P** |
| Intercept | | 0.7746 | | 0.11721 | | 6.609 | | 5.05E-10 |
| Red mean | | 0.34143 | | 0.28363 | | 1.204 | | 0.23 |
| Blue mean | | -0.09153 | | 0.12851 | | -0.712 | | 0.477 |
| Green mean | | -0.29047 | | 0.27448 | | -1.058 | | 0.291 |
| Residual deviance: 4601.1 on 166 degrees of freedom | | | | | | | | |
| dispersion parameter: 23.11532 | | | | | |  | |  |
| **Table S7. Pairwise contrasts between wing thickness for each wing panel, from a multi-response Bayesian mixed effects model** | | | | | | |  |  |
| **Contrast** | **Estimate** | | **Lower HPD** | | **Upper HPD** | |  |  |
| P1-P2 | -7.457 | | -8.31 | | -6.5631 | |  |  |
| P1-P3 | -2.324 | | -4.64 | | -0.0566 | |  |  |
| P1-P4 | -3.039 | | -4.56 | | -1.0759 | |  |  |
| P2-P3 | 5.172 | | 3.27 | | 7.0814 | |  |  |
| P2-P4 | 4.43 | | 3.25 | | 5.7344 | |  |  |
| P3-P4 | -0.739 | | -1.67 | | 0.1917 | |  |  |
